# Testing implications of the polygenic model for the genetic analysis of loci identified through genome-wide association

**DOI:** 10.1101/639955

**Authors:** Wenyu Zhang, R. Guy Reeves, Diethard Tautz

## Abstract

It has been proposed that many loci with no significant association in GWA studies can nonetheless contribute to the phenotype through modifier interactions with the core genes, implying a polygenic determination of quantitative traits. We have tested this hypothesis by using *Drosophila* pupal phenotypes. We identified candidate genes for pupal length determination in a GWA and show for disrupted versions of the genes that most are indeed involved in the phenotype, presumably forming a core pathway. We then randomly chose genes below the GWA threshold and found that three quarters of them had also an effect on the trait. We further tested the effects of these knockout lines on an independent behavioral pupal trait (pupation site choice) and found that a similar, but non-correlated fraction of them had a significant effect as well. Our data thus confirm the prediction that a large number of genes can influence independent quantitative traits.

**Impact statement:** Quantitative traits are similarly likely influenced by randomly picked loci as by loci identified in a genome-wide association study.

## Introduction

Organismal phenotypes, *i.e*., traits measured in whole organisms usually have a quantitative distribution and their genetic architecture can be studied by genome-wide association (GWA) approaches. In the past years, these approaches have suggested that such phenotypes have a polygenic architecture in the sense that many genes of moderate or small effect contribute to the phenotype (Wood et al., 2014; Yang et al., 2011, 2010). Hence, the attention has turned towards the loci falling below this cut-off. A study focussing on these small effect loci has suggested that all, or at least almost all, genes can be expected to contribute to the phenotype. This has led to the suggestion of an omnigenic model for quantitative traits (Boyle et al., 2017; Liu et al., 2019). It assumes that there could be a set of core genes forming pathways with special relevance for the phenotype, which are modified by many, if not all other genes expressed in the same cells or developmental stage. Although the effect sizes of these other genes are expected to be smaller than those of the core genes, in sum they explain more of the genetic variance or heritability than the set of core genes.

A general strategy to verify the genetic function of a locus that was identified through a GWA is to study the effect of an artificial knockout allele of the locus in an isogenic background. If this results in a phenotype that is related to the phenotype that was originally studied in the GWA, one considers the association as genetically confirmed. However, in a typical GWAS, one studies only naturally segregating alleles, which are usually not knockout alleles. Hence, one is asking very different genetic questions with these approaches, but these can potentially complement each other. A gene with an allele that segregates as a modifier of other pathways in natural populations may have a much stronger effect when it is disrupted by a major mutation, thus revealing its connections to the phenotype much more clearly. Intriguingly, under the omnigenic model, one would predict that basically every randomly chosen gene that is expressed in the relevant developmental stage could show an effect on the phenotype studied, when it is disrupted by a major mutation. We have tested this hypothesis here, by comparing the genetic effects of gene disruption lines for loci that were identified through a GWA versus loci that were randomly chosen.

We have chosen *Drosophila* pupal size and pupation site choice as quantitative phenotypes, for which we have previously developed automated phenotyping pipelines and have shown their polygenic basis (Reeves and Tautz, 2017; Zhang et al., 2020). The pupal stage is an immobile life history stage of transformation between larval and adult stages found in holometabolous insects (Jones and Reiter, 1975; Price, 1970). Pupal size (or pupal case length) can be considered as an analogue to human height, in terms of morphological measure and various biological properties, such as equal paternal and maternal impact on offspring trait variance (Reeves and Tautz, 2017). Pupation site choice (or pupation height) is measured as the distance of the pupation site above the surface of the food. It reflects an ecologically relevant trait that is behaviourally determined and where a core network could be identified (Zhang et al., 2020). The phenotypes are not directly related to each other (Zhang et al., 2020).

For assessing the genetic contribution to these quantitative traits, we use the extensive genetic resources available for *Drosophila*. This includes the *Drosophila* Genetic Reference Panel (DGRP) as a resource for association mapping, as well as the gene disruption lines maintained at the Bloomington *Drosophila* Stock Centre (BDSC). The DGRP consists of a population of more than 200 fully sequenced highly inbred *Drosophila melanogaster* strains derived from a wild population in the Raleigh, US (Huang et al., 2014; MacKay et al., 2012), which has been successfully applied to detect the association between a broad range of complex traits and their underlying genetic basis (Dembeck et al., 2015; Durham et al., 2014; Huang et al., 2014). Together with the well-resolved richness in genetic polymorphisms and rapid decay in linkage disequilibrium (LD) in these strains (Huang et al., 2014), these make the DGRP an excellent resource to study the genetic architecture of pupal phenotypes in *Drosophila melanogaster* at a fine scale resolution. The BDSC gene disruption lines were created through systematic transposon mutagenesis experiments (Bellen et al., 2011). They can be used to test GWA candidate genes for a possible involvement in the trait, but also as a source for randomly chosen genes to study their effect on a quantitative trait.

In the present study, we conduct a standard GWA for pupal size based on the DGRP panel and use gene expression profiling analysis to reveal the relevant developmental stages for this trait. Following the predictions of the omnigenic model, we explore also the effects of genes randomly chosen among genes expressed in the respective life stage and with no significant GWA score. We find that not only a large fraction of the GWA predicted genes, but also of the randomly chosen genes have an effect on the pupal case length phenotype. Moreover, we find that many of the same genes affect also the pupation site choice phenotype, although the effect sizes between these two phenotypes are not correlated. We conclude that these findings support predictions of an omnigenic genetic architecture for quantitative traits and proof that association studies versus classic genetic studies reveal indeed different aspects of the genetic architecture.

## Results

### Phenotyping

We used pupal case length, defined as the major axis length in millimetre (mm) of the pupa as an ellipse-shape object (Figure 1), as a measure of pupal size in *Drosophila melanogaster.* We adapted and modified the earlier image-analysis based phenotyping pipeline (Reeves and Tautz, 2017) for the high-throughput measurement of pupal case length. The same approach allows also to score the independent phenotype of pupation site choice (pupation height) and for both we have shown that a very large number of phenotypes can be measured with high precision (Reeves and Tautz, 2017; Zhang et al., 2020).

**Figure 1:**
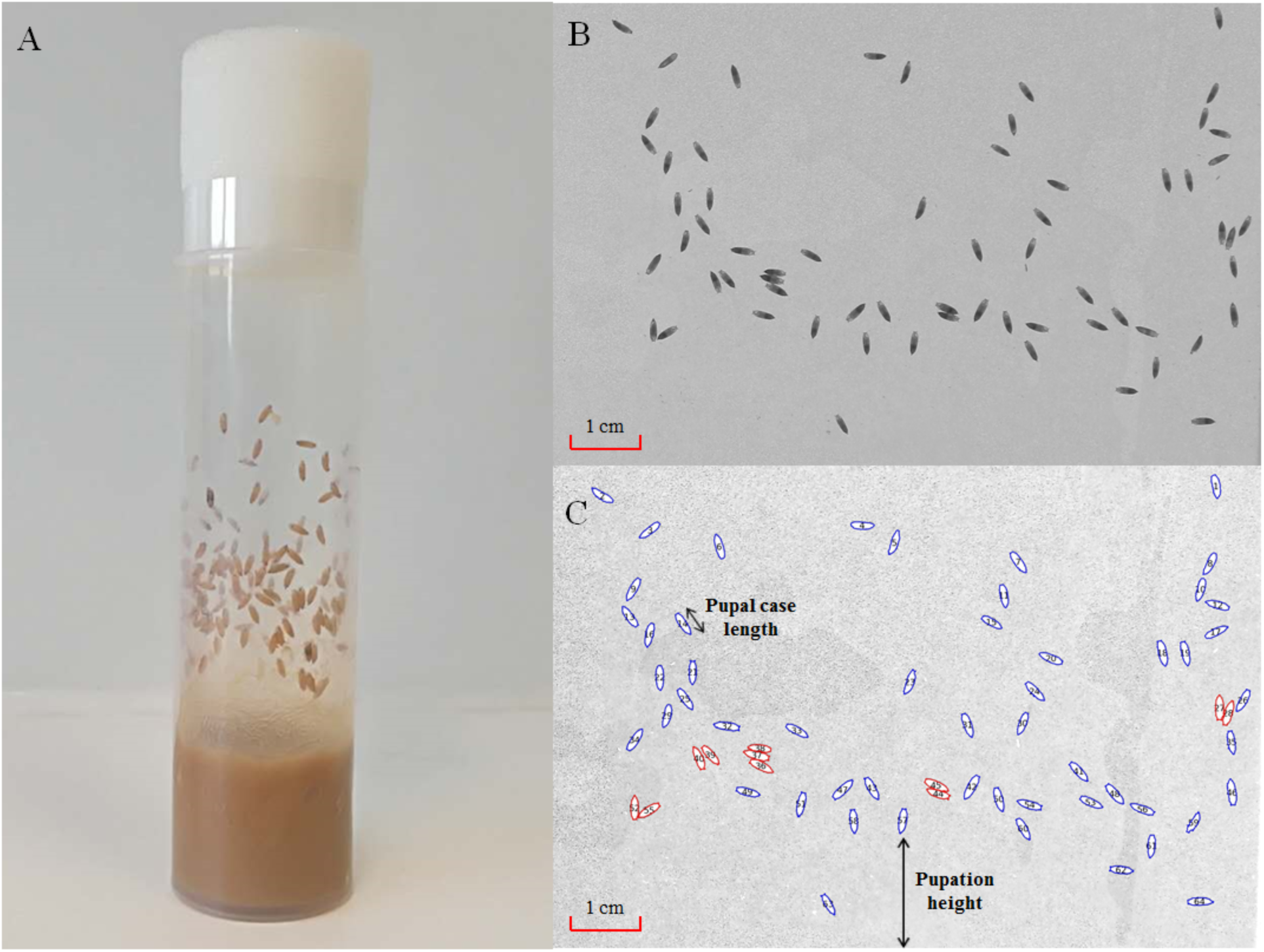
Automated measurement of pupal case length and pupation height. (A) shows a vial with sufficient number of pupae attached in the wall. A plastic sheet that lines the wall is included inside the vial, and is ready to be taken out for being photographed (B). The image analysis software CellProfiler (Carpenter et al., 2006) is then used to identify the outlines of the pupae and measure their major axis lengths and distances between their pupation sites to the food surface (C). The pupae that can be reliably measured (singularized) are marked in blue, and the ones with lower confidence scores (aggregated) are marked in red.

To assess whether the DGRP lines reflect the scope of natural variation, we measured also the pupal case length phenotypes of 14 natural wild-type *Drosophila melanogaster* strains collected from 9 global geographic regions (strain details in Supplementary Table S1a and S1b). Two of these wildtype stocks (S-317 and S-314) were continually re-measured to act as controls, and showed consistent measurement on pupal case length throughout all experiments (Figure 2–figure supplement 1).

The profiles of pupal case length from the wildtype and of 198 inbred lines from the *Drosophila* Genetic Reference Panel (DGRP) (Huang et al., 2014; MacKay et al., 2012) are shown in Figure 2. We observed a large variation of average pupal case length among strains, ranging from 2.9 mm to 3.4 mm for natural wild-type strains, and ranging from 2.7 mm to 3.5 mm for DGRP inbred stocks. The spread of pupal case length among the DGRP inbred lines exceeded that of the DSPR recombinant inbred stocks (ranging from 2.9 mm to 3.5 mm) (Reeves and Tautz, 2017), suggesting that DGRP lines capture indeed the existing variation of pupal case length in *Drosophila melanogaster*. The same conclusion applies also to the pupation site choice phenotype, as we have shown previously (Zhang et al., 2020).

**Figure 2:**
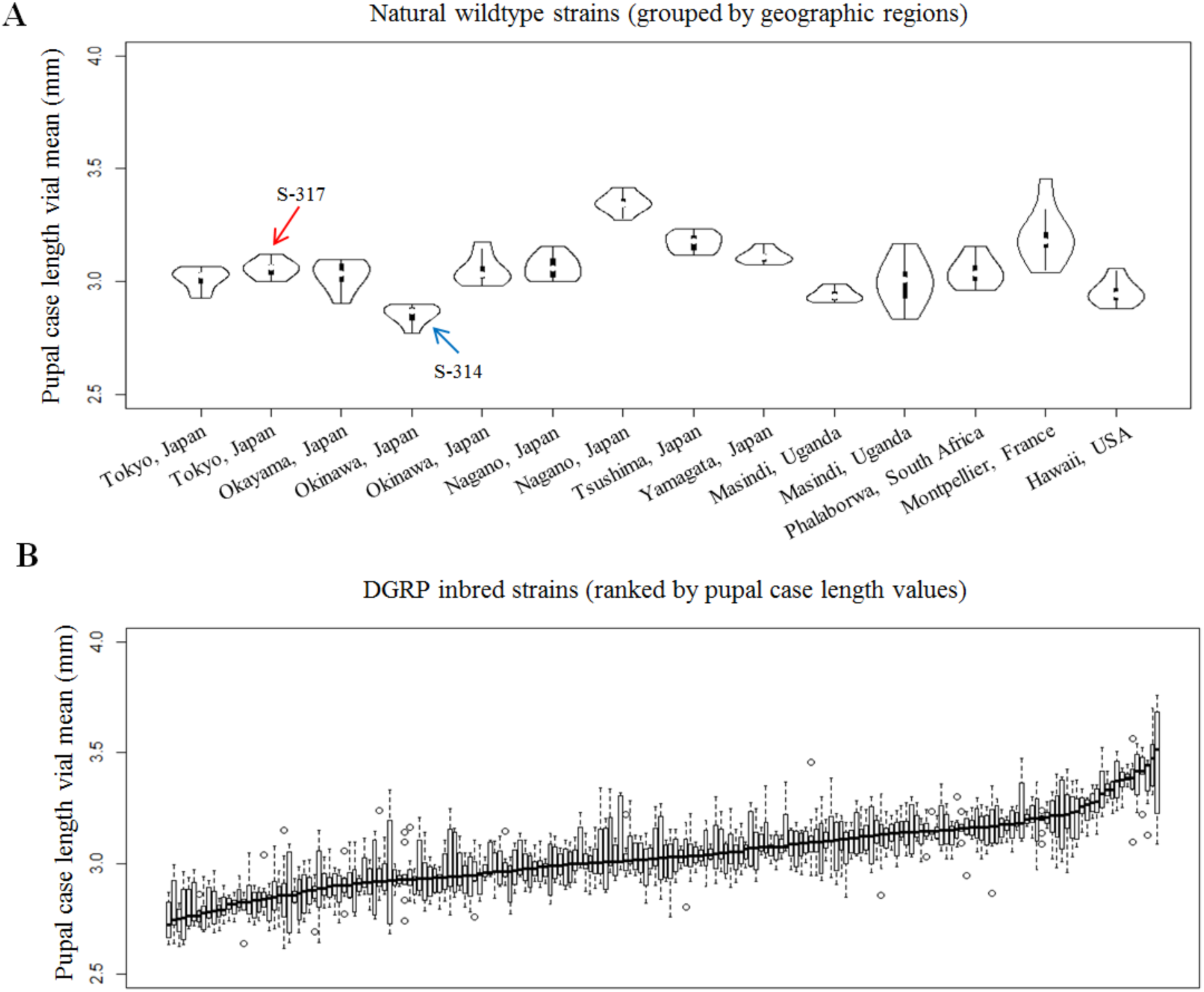
Pupal case length distribution for different *Drosophila melanogaster* strains. The distribution of natural wild derived stocks is grouped by geographic regions (A), and that of DGRP inbred strains is ranked by pupal case length values. The two natural stocks (S-317 and S-314) acting as controls are labelled in the panel (A). **Figure 2–figure supplement 1:** Repeated measurement of pupal case length of two control stocks.

### Heritability and chromosome effects

The estimate of the total genetic component of pupal case length, *i.e.*, broad sense heritability (H^2^), was determined as the proportion of total variance in the mean strain pupal case length measurements compared to the average within each strain (Schmidt et al., 2017). Our estimates of values of H^2^ (0.71 for wild-type strains; 0.52 for DGRP inbred strains) were similar to the ones analysed with the same methodology, but based on another set of *Drosophila melanogaster* strains (Table 1) (Reeves and Tautz, 2017). Combining with the previous estimates on narrow sense heritability (h^2^, ranging from 0.44-0.50, Table 1) (Reeves and Tautz, 2017), our analysis suggested that the additive genetic impact (*i.e.*, narrow sense heritability) contributed to a major part (∼78%) of total genetic component of pupal case length in *Drosophila melanogaster.*

**Table 1.** Statistics for the heritability estimates of pupal case length.

| Heritability | Method | Dataset | Heritability Estimate |
| --- | --- | --- | --- |
| $H^2$ | IBM SPSS<br>(Variance component) | <sup>a</sup>wild-type strains | 0.71 |
|  |  | <sup>b</sup>DGRP inbred strains | 0.52 |
|  |  | <sup>c</sup>Four-Way | 0.58 |
|  |  | <sup>d</sup>Eight-Way (DSPR) | 0.61 |
| $h^2$ | Mid-parent regression | <sup>e</sup>Four-Way | 0.44 ± 0.04 SE |
|  |  | <sup>f</sup>Eight-Way (DSPR) | 0.50 ± 0.09 SE |

<sup>a</sup> Number of stocks = 14; Average number of replicates per stock = $7.8 \pm 0.6$ SD; Average number of measured pupae per vial = $69 \pm 24$ SD;
<sup>b</sup> Number of ILs = 198; Average number of replicates per stock = $8.2 \pm 1.6$ SD; Average number of measured pupae per vial = $40 \pm 17$ SD;
<sup>c</sup> Number of RILs = 81; Average number of replicates per stock = $5.5 \pm 0.8$ SD; Average number of measured pupae per vial = $59 \pm 24$ SD;
<sup>d</sup> Number of RILs = 195; Average number of replicates per stock = $6.7 \pm 2.4$ SD; Average number of measured pupae per vial = $70 \pm 34$ SD;
<sup>e</sup> Number of single-pair crosses = 363; Average number of measured pupae per vial = $62 \pm 17$ SD;
<sup>f</sup> Number of single-pair crosses = 67; Average number of measured pupae per vial = $47 \pm 15$ SD;
<sup>c-f</sup> Datasets were retrieved from (Reeves and Tautz, 2017);
ILs: Inbred lines; RILs: recombinant inbred lines; DSPR: *Drosophila* Synthetic Population Resource;
SD: standard deviation; SE: standard error;

We partitioned the explained phenotypic variance by chromosome, via calculating the “SNP heritability” in the DGRP stocks (Wray et al., 2013), *i.e.*, the estimate of the proportion of phenotypic variance explained by all available SNPs (or genetic variants) within each chromosome. Our results show a much higher contribution of autosomes to the variance of pupal case length (Table 1-figure supplement 1), with the exception of chromosome 4, mostly likely due to the limited number of genetic variants within this chromosome. Hence, pupal case length is a highly heritable phenotype and the DGRP lines include the genetic variation suitable for mapping, similarly as we have previously shown for the pupation site choice phenotype (Zhang et al., 2020).

### Genome-wide association analysis

We used the genetic variants of DGRP freeze 2 (Huang et al., 2014), and excluded the variants with missing values above 20% and minor allele frequency below 5% from the further analysis. Several possible covariates were assessed, in order to optimize the model for GWA mapping analysis (Figure 3).

**Figure 3:**
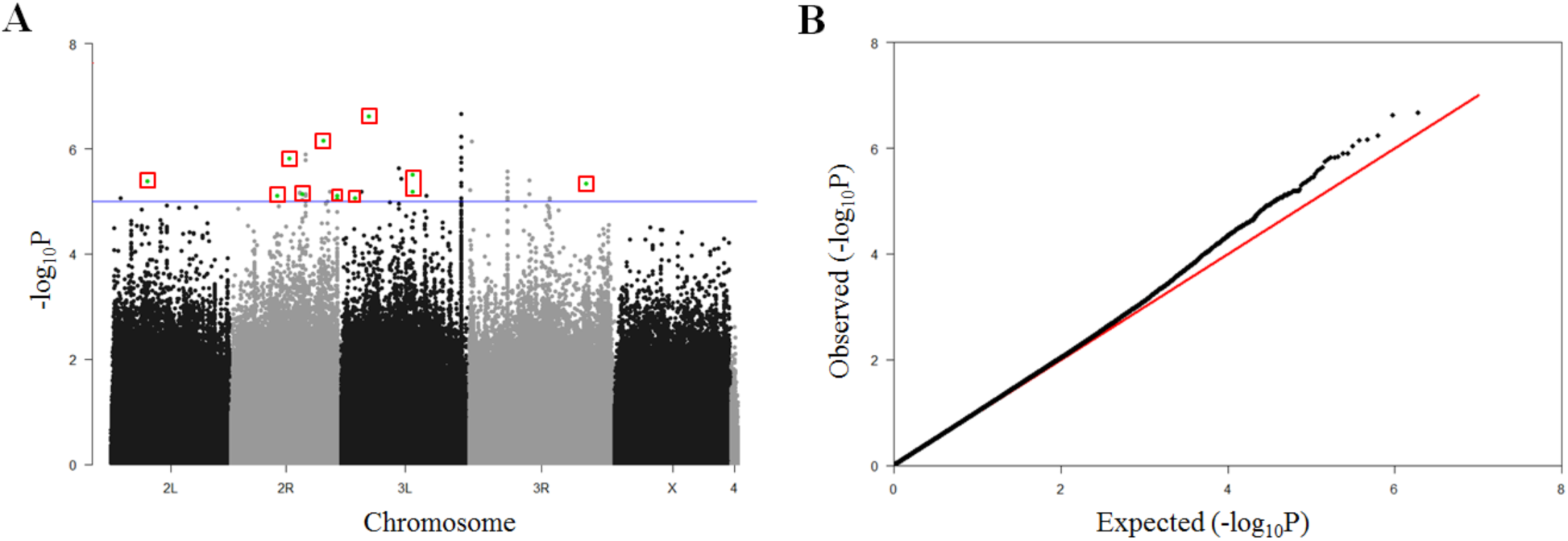
GWA mapping results on pupal case length. (A) Manhattan plot for GWA results. The p-values in −log10 transformation are shown on the y-axis. The blue horizontal line marks the nominal p-value threshold (1 × 10^−5^) used in this study. The genetic variants in red rectangles are the ones selected for experimental validation as shown in Figure 4. (B) Quantile-quantile (Q-Q) plot for GWA results. The expectation and observation of GWA p-values in −log_10_ transformation are represented by the x-axis and y-axis, respectively. **Figure 3-figure supplement 1:** Effect of *Wolbachia* infection on pupal case length. **Figure 3-figure supplement 2:** Correlation between pupal case length and population structure. **Figure 3-figure supplement 3:** Correlation between pupal case length and top 20 PCs. **Figure 3-table supplement 1:** Comparison of pupal case length between Wolbachia-free and Wolbachia-infected stocks. **Figure 3-table supplement 2:** Correlation between pupal case length and inversion status in DGRP strains.

Around half of the DGRP strains are infected by *Wolbachia pipientis*, a maternally transmitted endosymbiotic bacterium (Huang et al., 2014) (Supplementary Table S1b). We examined the possible effects of *Wolbachia* infection on pupal case length through two different approaches. We first compared the pupal case length values between infected strains and non-infected strains, and found no significant difference between these two groups (Wilcoxon rank-sum test: p-value=0.90, (Figure 3- table supplement 1). Second, we randomly selected three DGRP strains with *Wolbachia* infection, and tested the phenotypic change on pupal case length after the removal of *Wolbachia* infection by using tetracycline treatment. No significant statistical differences on pupal case length were observed for all tested strains (Figure 3-figure supplement 1). Based on these analyses, we concluded that the *Wolbachia* infection on DGRP strains had only minor, if any, influence on pupal case length, and consequently we did not incorporate the *Wolbachia* infection status in the association analysis below.

The DGRP lines were generated in a way that population structure impacts should be minimized, but some genetic relatedness leading to cryptic population structure might still exist (Mackay and Huang, 2018). To examine whether any cryptic population structure could contribute to the observed pupal case length variation of DGRP inbred stocks, we exploited PLINK (Purcell et al., 2007) to identify major principal components (PCs) of genetic variants in the DGRP strains. Based on the projection analysis of the top 2 PCs, we found no obvious clusters on the basis of pupal case length value classes (Figure 3-figure supplement 2). However, further correlation analysis revealed three out of the top 20 PCs to be significantly associated with pupal case length values (Figure 3-figure supplement 3, Pearson’s correlation test, p-value⩽0.05), suggesting a possible impact on pupal case length from cryptic population structure. In order to correct any potential influence from the cryptic population structure, we decided to use a linear mixed model implemented in fastlMM (Lippert et al., 2011) program (version 0.2.32) for the GWA mapping analysis.

Approximately 45% of the DGRP strains harbour at least one type of major genomic inversion (Huang et al., 2014) (Supplementary Table S1b), and these major genomic inversions might contribute to the observed population structure and have impact on pupal case length as well. We systematically tested the correlations between genomic inversion status in DGRP strains and the top 2 PCs based on genetic variants as mentioned above, and found significant effects from In(2L)t and In(3R)Mo (Pearson’s correlation test, p-value⩽0.05, Figure 3-table supplement 2), hinting their potential roles in population divergence (Hoffmann and Rieseberg, 2008). Moreover, we also observed a significant association between pupal case length and In(3R)Mo (Pearson’s correlation test, p-value=0.0002). Accordingly, the presence status of In(3R)Mo in DGRP lines was incorporated as a covariate of the linear mixed model used for GWA mapping analysis on pupal case length. We used a significance cut-off of p-value < 1 × 10^−5^, which is a nominal threshold frequently used in *Drosophila* quantitative trait genetic studies (Dembeck et al., 2015; Lee et al., 2017; Zhang et al., 2020). The mapping revealed a number of associations above the chosen threshold (Figure 3A) and the qq-plot shows that there is substantial genetic variation above the random expectation (Figure 3B).

### Candidate genes from the GWA analysis

We found 50 significant SNPs to be associated with pupal case length in the DGRP strains, corresponding to 67 associating genes that locate within 5kb up/down-stream (default setting in SnpEff (Cingolani et al., 2012)) of these genetic variants (Table 2). The effect sizes of individual SNPs range from 0.5% to 3.3% of the total phenotypic variation (Table 2-figure supplement 1).

**Table 2:**
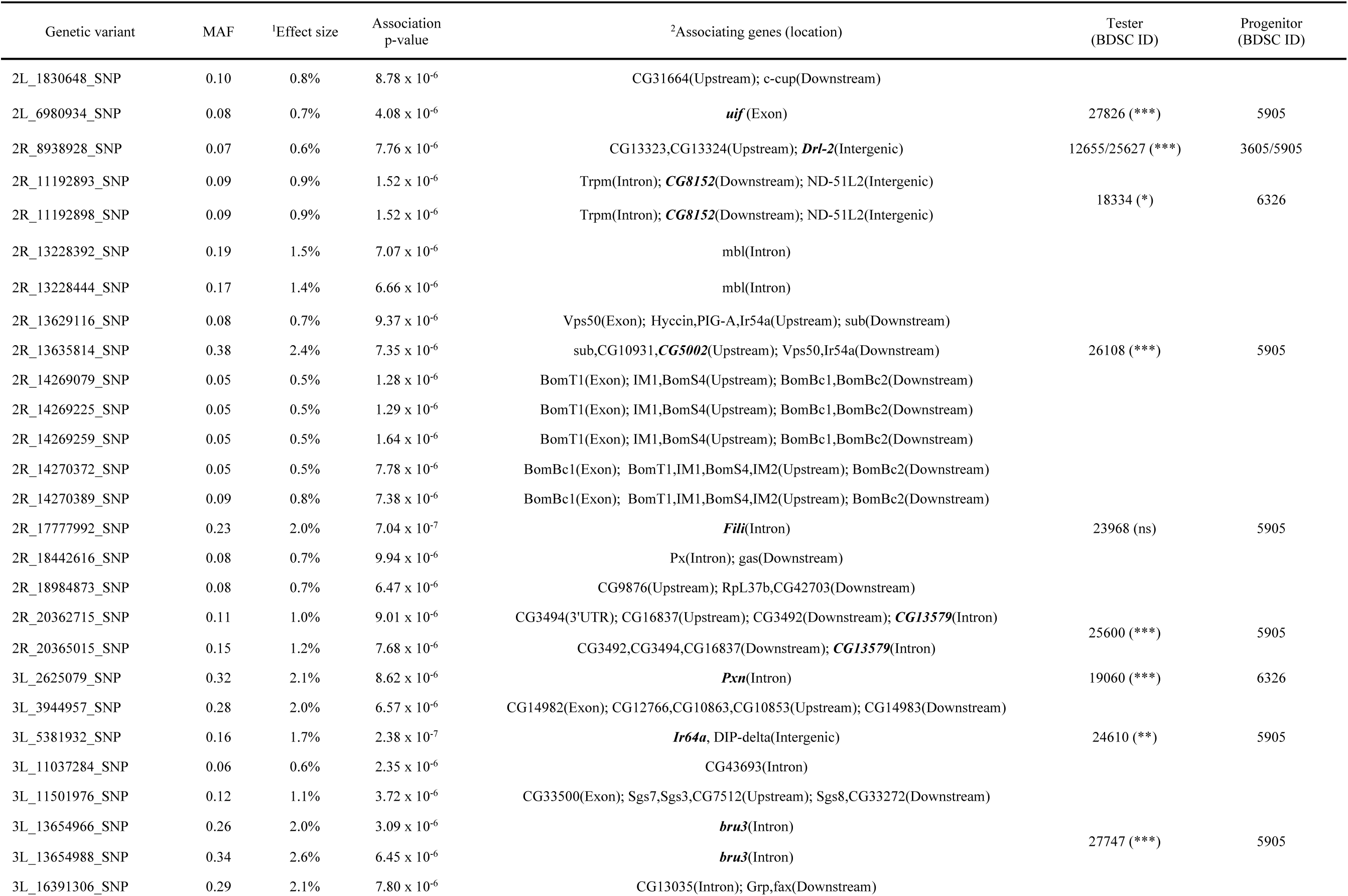
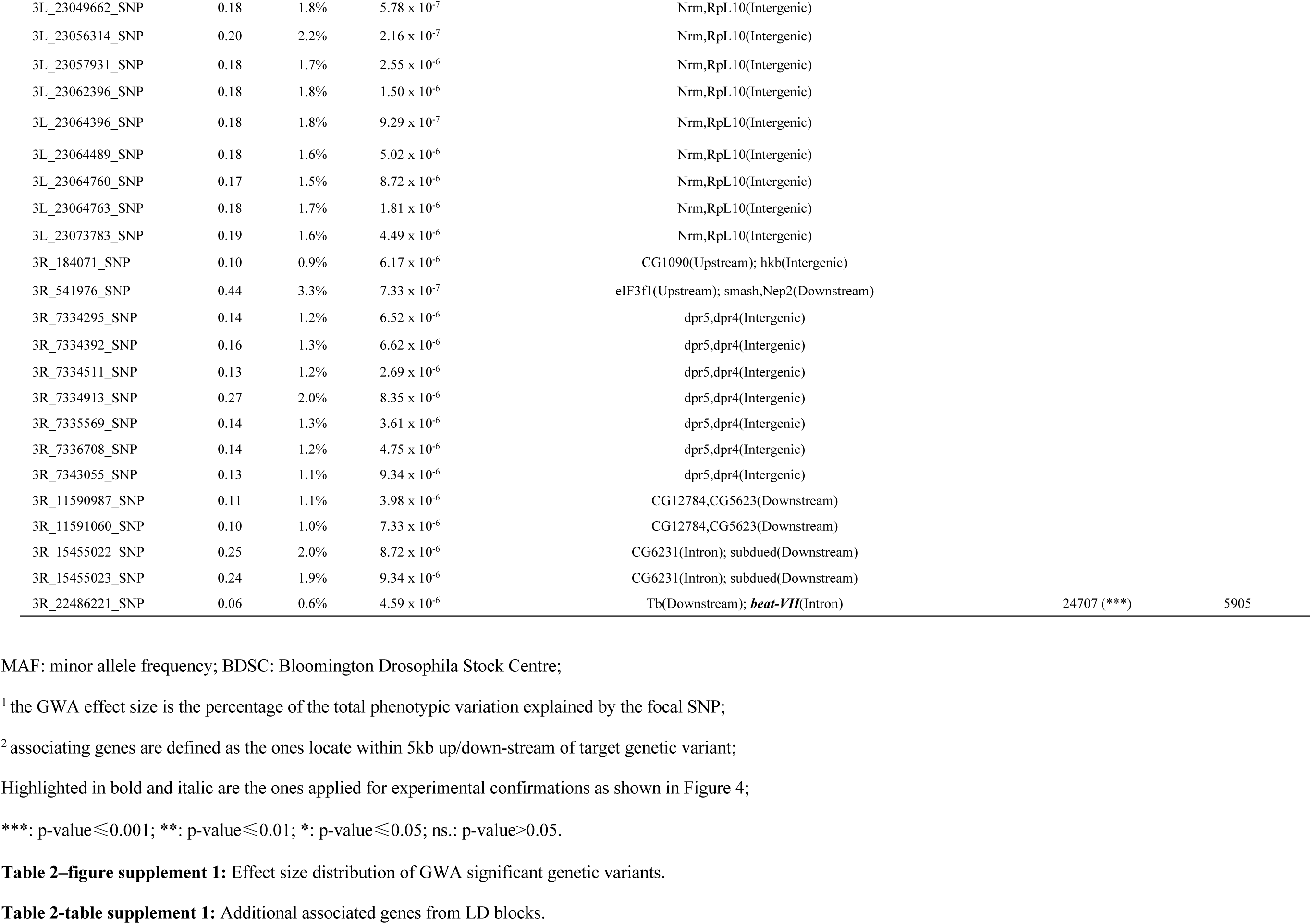
Genome-wide association results on pupal case length.

To identify possible additional candidate genes associated with the variants, we examined the long-range linkage disequilibrium (LD) between pairs of detected candidate variants and with other genetic variants found in the DGRP strains. LD blocks were then calculated for each significant genetic variant with a commonly used threshold r^2^ = 0.8 (Pallares et al., 2014), and 17 significant LD blocks were found with average block size of 19.3 kb (Table 2-table supplement 1). This finding is in line with the observation of a rapid decay of LD in the panel strains reported in the original DGRP resource reference (Huang et al., 2014). Combining the additional genes identified in the above LD blocks, we identified in total 90 candidate genes associating with pupal case length variation in *Drosophila melanogaster*.

### Phenotype confirmations

To test the phenotypic effects from different alleles on pupal case length at single gene scale, we used transposon insertion mutagenesis lines that have been constructed in a common co-isogenic background (Bellen et al., 2011). Eleven gene disruption lines corresponding to 10 of the GWA identified associated genes were available for this experiment (Table 2). All experimental tests were done via replicated phenotypic comparisons between co-isogenic stocks and homozygous mutant lines, except for the gene of *Pxn*, which was tested as heterozygote (Figure 4), as the homozygous disruption of this gene is lethal (see Methods).

**Figure 4:**
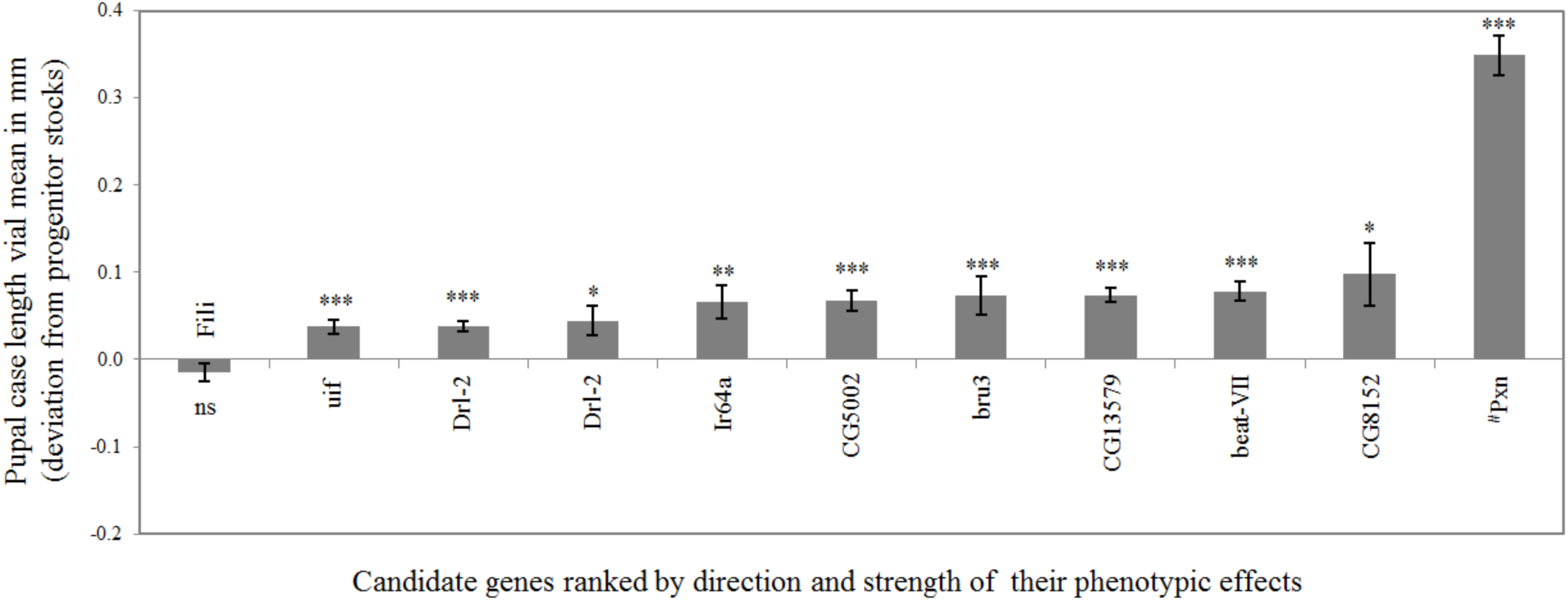
Gene disruption test results on the pupal case length phenotypic effects of GWA candidate genes. Phenotypic effects of candidate genes were from transposon insertion tests. The phenotypic effects were measured as the deviation of pupal case length of stocks with gene disruption compared with that from corresponding progenitor stocks. The error bars show the standard error of mean (SEM) values. Statistical p-values were computed via Wilcoxon rank-sum tests. ***: p-value⩽0.001; **: p-value⩽0.01; *: p-value⩽0.05; ns: p-value>0.05. Stock marked with “^#^” was tested as heterozygotes; the remaining were stocks tested as homozygotes.

Figure 4 shows the profile on the measured pupal case length differences between mutant lines and the respective progenitor stocks. All apart one of the lines showed a significant difference (Wilcoxon rank-sum test, p-value 0.05), interestingly all towards an increased pupal case length, implying that the effect is not just due to a general physiological disruption of pupal growth. No significant correlation was found between GWA predicted effect sizes and the phenotypic effect sizes from the above gene disruption tests (Pearson’s correlation test, r value = 0.28, p-value = 0.44), confirming the notion that naturally segregating alleles have usually different genetic effect sizes when compared to disruption alleles. It should be noted that two independent experimental tests were conducted for gene *Drl-2*, with transposon insertions landing in the different regulatory regions and from different co-isogenic backgrounds (Supplementary Table S1c). Both showed consistent phenotypic effects, affirming the robustness of the gene disruption assays.

### Expression analysis

Genes involved in determining pupal case length are expected to be expressed at developmental stages close to the pupation process. To explore this, we analysed a developmental RNA-Seq dataset from *Drosophila melanogaster* (Graveley et al., 2011), which included the transcriptome of 27 distinct developmental stages covering all the major phases of the *Drosophila* life cycle. For the ease of presentation, we further grouped these 27 distinct developmental stages into six super-developmental stages: early embryo, middle embryo, late embryo, larval stage, pupal stage, and adult stage (Figure 5).

**Figure 5:**
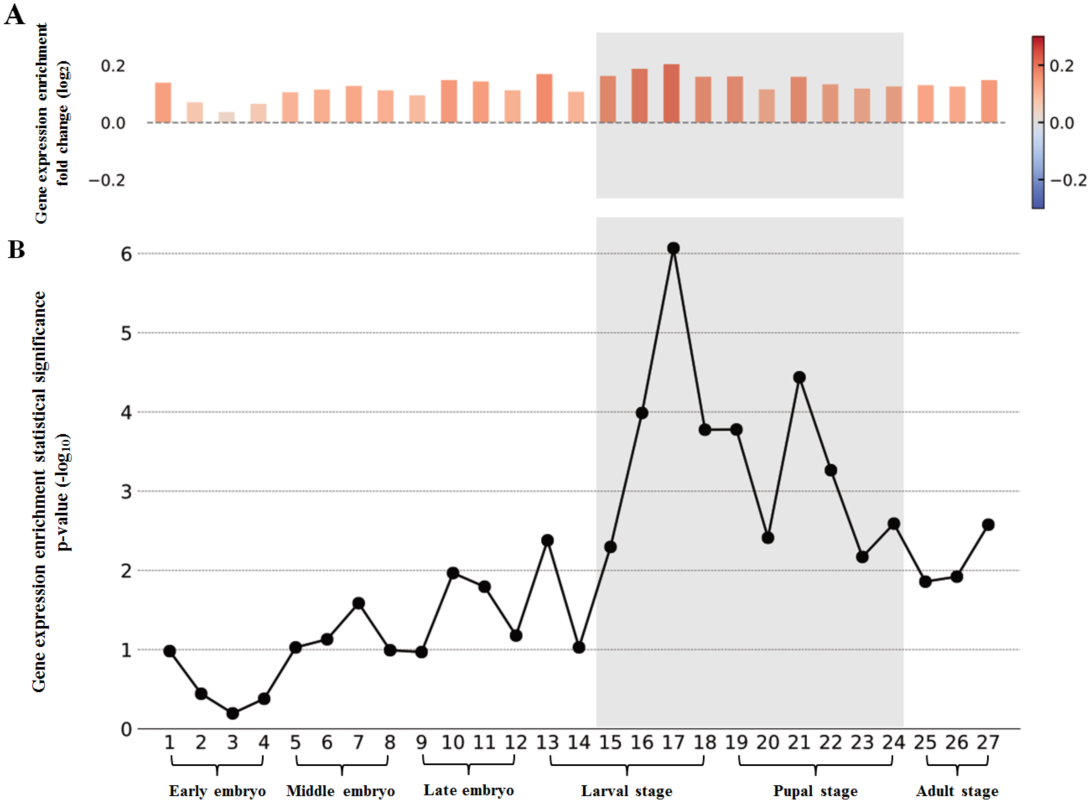
Gene expression enrichment patterns of GWA candidate genes across different developmental stages. The expression enrichment statistics of GWA candidate genes were calculated with total annotated protein coding genes as a control. The panel (A) shows the gene expression enrichment fold changes (*i.e.*, the ratios between the fractions of expressed GWA candidate genes and total coding genes, in log2 scale) between GWA candidate genes and total coding genes, and the panel (B) shows the statistical significances (p-values in –log10 scale, Fisher’s exact test) of the above gene expression enrichment patterns in 27 developmental stages. *Early embryo*: 1) 0-2 hrs. embryo; 2) 2-4 hrs. embryo; 3) 4-6 hrs. embryo; 4) 6-8 hrs. embryo; *Middle embryo*: 5) 8-10 hrs. embryo; 6) 10-12 hrs. embryo; 7) 12-14 hrs. embryo; 8) 14-16 hrs. embryo; *Late embryo*: 9) 16-18 hrs. embryo; 10) 18-20 hrs. embryo; 11) 20-22 hrs. embryo; 12) 22-24 hrs. embryo; *Larval stage:* 13) L1 stage; 14) L2 stage; 15) L3 stage of 12 hr. post-molt; 16) L3 stage of dark blue gut (PS1-2); 17) L3 stage of light blue gut (PS3-6); 18) L3 stage of clear blue gut (PS7-9); *Pupal stage:* 19) white prepupa (WPP), 20) 12 hr. after WPP (P5), 21) 24 hr. after WPP (P6); 22) 2 days after WPP (P8), 23) 3 days after WPP (P9-10), 24) 4 days after WPP (P15); *Adult stage:* 25) male/female after 1-day eclosion; 26) male/female after 5-day eclosion; 27) male/female after 30-day eclosion. For adult developmental stages, the enrichment results for male and female were averaged for presentation. The shaded area indicates the “relevant” developmental stages of pupal case length morphogenesis. **Figure 5-table supplement 1:** Summary on gene expression patterns across different developmental stages.

Compared with total annotated coding genes as a control, we observed an enrichment of GWA candidate genes with expression (FPKM >0) in all 27 tested distinct developmental stages (Figure 5A and Figure 5-table supplement 1), suggesting that GWA tends to pick up gene candidates with broad expression. More intriguingly, we found these GWA candidate genes are mostly significantly enriched in the developmental period between late larval stage and pupal stage (Fisher’s exact test, p-value0.01, Figure 5B). As the pupation process occurs from the late larval stage, and the genesis of pupal case is completed before eclosion, our results imply that the developmental period between late larval and pupal stage might be the most relevant developmental stages for pupal case length morphogenesis in *Drosophila melanogaster*.

### Phenotypic effects of randomly chosen genes

The omnigenic model (Boyle et al., 2017) predicts that most, if not all genes can affect a given quantitative trait, when they are expressed in the relevant developmental stage. We have therefore set out to use our phenotyping pipeline to test this prediction. We selected 47 gene disruption lines (corresponding to 45 disrupted genes, all tested as homozygotes) constructed in a co-isogenic background from the panel of *Drosophila* gene disruption stocks (Bellen et al., 2011) using the following criteria: 1) the GWA p-value of the focal gene falls below the significance threshold (*i.e.*, > 1 × 10^−5^); 2) the corresponding gene has detectable expression in the above defined relevant developmental stages for pupal case length (FPKM >0), and 3) homozygous disruption is viable. The latter criterion biases against essential genes (approximately 30% are not homozygous viable).

Figure 6A compares the phenotypic effect sizes of all strains tested in this study with their respective pupal case length GWAS p-values. It shows that the genes picked because of their GWA significance have not necessarily the largest phenotypic effects as disruption lines, and that there is no significant difference on the overall phenotypic effects between GWA candidate genes and randomly chosen genes (Wilcoxon rank-sum test p-value=0.39, Figure 6-figure supplement 1A). The conclusion remained the same when the only one GWA predicted gene that was tested as heterozygotes (*Pxn*, Figure 4) was removed from the analysis (Wilcoxon rank-sum test p-value=0.14). Overall, around 76% of random genes (34 of 45) showed significant effects (p-value ≤ 0.05) on the pupal case length phenotype (Figure 6B), in contrast with the 90% of GWA candidate genes (9 out of 10), but these fractions are not significantly different (p-value = 0.43, Fisher’s exact test).

**Figure 6:**
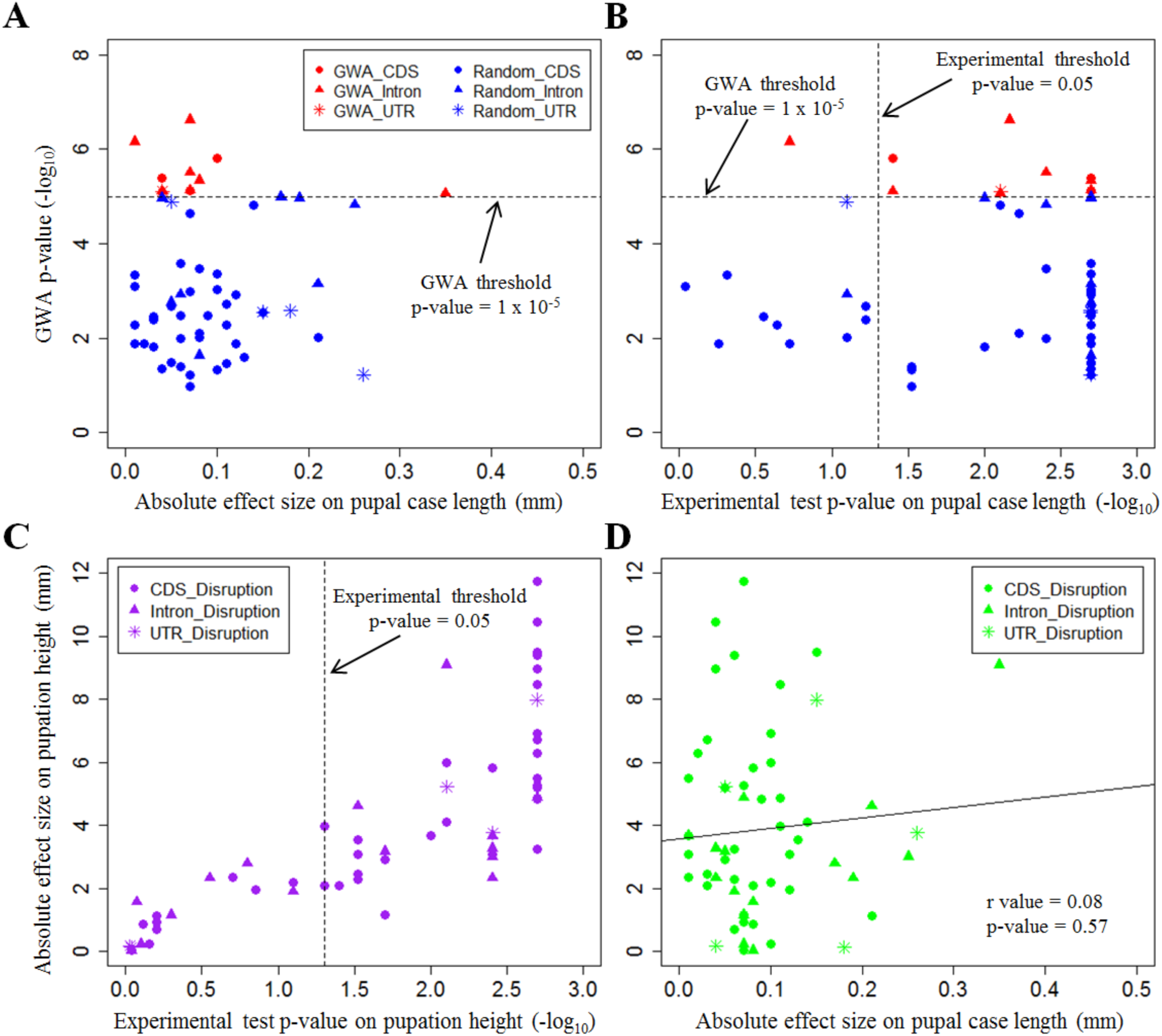
The impact of gene disruption on pupal case length and pupation height. Comparison between GWA p-values and absolute experimental effect sizes and statistical p-values for gene disruption of GWA candidate genes (labelled in red) and randomly selected genes (labelled in blue) on pupal case length are shown in (A) and (B), respectively. The GWA p-value is defined as the lowest p-value from all the genetic variants within the 5kb up/down-stream of target gene (including the gene itself). (C) Pupation site choice (height) comparison between absolute experimental effect sizes and statistical p-values for gene disruption of all tested genes. Experimental statistical p-values were computed via Wilcoxon rank-sum tests. The horizontal dashed line marks the GWA nominal p-value threshold (1 × 10^−5^), and the vertical dashed line indicates the p-value threshold for experimental tests (0.05). (D) Correlation between the absolute experimental effect sizes on pupation height and pupal case length. The statistics on the correlation was computed by using Pearson’s correlation test. Each dot represents one gene disruption line. Genes with transposon insertion mutagenesis in different genic regions (CDS, Intron, or UTR) are labelled with various dot shapes. The absolute experimental effect size was measured as the absolute deviation of pupal case length/pupation height of the stock with the target gene disruption compared with that from corresponding progenitor stock. **Figure 6-figure supplement 1:** Gene disruption effect sizes on pupal case length based on their categories. **Figure 6-figure supplement 2:** Genetic variant density comparison between GWA genes and random genes with significant pupal case length phenotypic effects. **Figure 6-table supplement 1:** Detailed information on all the tested gene disruption lines.

It should also be noted that these strains may 1) have different co-isogenic backgrounds, 2) have disruption in different regions, either coding regions (disruption on the structure of coded proteins) or regulatory regions (alteration of the gene expression levels). However, these two factors seemed to play only minor roles in the phenotype differences, given the observation that similar extents phenotypic effects on pupal case length can be observed for the disruption of the same gene from difference genetic background and/or different gene regions (*Drl-2, CG14007*, and *CG42260*. See Figure 6-table supplement 1). Moreover, systematic comparison on the effect size on pupal case length also showed no significant difference either on the disruption of different gene regions (Figure 6-figure supplement 1B), or the percentage of disruption on the gene’s CDS regions (Figure 6-figure supplement 1C).

### Phenotypic effects on an independent phenotype

If most genes have an effect on one quantitative trait, one would expect that the same is true for a second trait. As discussed above, pupation site choice is such a second independent trait, that we measured with the same setup. Given the sharing of gene expressions in the common relevant developmental stages (Figure 6-table supplement 1), all the disrupted genes (both the 10 tested GWA genes and 45 random genes) on the basis of pupal case length can therefore serve as random gene disruption tests for pupation site choice. Indeed, we observed that around 67% of the randomly tested genes (38 of 55) actually showed significant effects (p-value⩽0.05) on the pupation site choice trait (Figure 6C). This fraction of effects is not significantly different from the fraction of effects of randomly tested genes for the pupal case length phenotype reported above (p-value = 0.51, Fisher’s exact test). This supports the notion that the majority of genes with expression in the relevant developmental stages can have effects on a given quantitative trait at this stage. Interestingly, the effect sizes do not correlate for the two phenotypes tested (Figure 6D), implying that the effects are channelled through different networks.

## Discussion

We have here tested predictions of the omnigenic model for quantitative traits, especially the question of the possible role of modifier loci without significant GWA score (Boyle et al., 2017; Liu et al., 2019). Although it is debated whether the term “omnigenic” is more useful than the long established terms “polygenic” and “infinitesimal” (Wray et al., 2018), the analysis by (Boyle et al., 2017) has certainly sparked new interest in this almost century-old question. But we are only now coming into a phase where predictions from these models can be directly tested. Most of the evidence has so far come from human studies, which are often focussed on disease questions and their associated special considerations and limitations (Wray et al., 2018). But for well-developed genetic model systems, such as *Drosophila*, one can do direct genetic experiments.

The *Drosophila* system allows to systematically using both, a GWA approach, as well as the classic knock-out approach to assess the effect of loci on given phenotypes. Furthermore, the availability of automated phenotyping pipelines allows to score a large number of individuals under highly controlled conditions and thus get a statistically sound resolution even for small effects. We have explored this here for two pupal phenotypes that are particularly amenable to analysis through automatic image processing.

Using the DGRP lines that represent segregating wildtype alleles and the commonly used corrections for confounding factors, we were able to identify a set of 90 candidate genes within 50 associated genetic loci for the pupal size phenotype (Table 2 and Table 2-table supplement 1). As expected for a typical polygenic trait, each genetic variant could only explain a small fraction (< 4%) of total phenotypic variance, implying that most of the heritability of the trait comes from a large number of loci with GWA scores below the chosen threshold. Using the transposon insertion mutagenesis lines that have been constructed in a common co-isogenic background (Bellen et al., 2011), we found that nine out of ten tested GWA candidate genes showed significant effects (p-value⩽0.05) on pupal size. These include the well-studied gene *Tubby (Tb)*, which was reported to be involved in body morphogenesis. *Tubby* is actually also a phenotypic pupal marker for a balancer chromosome (Lattao et al., 2011). Other clear candidate genes include two loci known to be involved in developmental processes, such as *uninflatable* (*uif)*, which modulates Notch signaling (Loubéry et al., 2014) and the receptor tyrosine kinase *Derailed 2* (*Drl-2*) that transduces Wnt5 signaling (Sakurai et al., 2009). Accordingly, the GWA did identify loci that could be part of core pathways of the phenotype, similar as we showed it for the pupation site choice phenotype (Zhang et al., 2020).

Gene expression analysis on GWA candidate genes revealed their enrichment in the developmental interval from late larval to the pupal stage, suggesting these developmental stages to be relevant of pupal case length phenotype. The omnigenic model (Boyle et al., 2017) predicts that most, if not all genes can affect a given quantitative trait, when they are expressed in the relevant developmental stage and organ(s). We thus set out to use gene disruption experimental tests on random genes with expression in the relevant developmental stages. The GWA p-value for these 45 randomly chosen genes is below any threshold that one would normally consider using. Accordingly, none of them would have been identified as GWA candidate genes. However, our results showed that approximately three quarters of them actually had significant effects (p-value⩽0.05) on the pupal case length phenotype. Intriguingly, a similar fraction, but only partially overlapping loci, showed also an effect on the behavioural pupation site choice phenotype.

One can ask why these randomly chosen genes, especially the ones showing significant phenotypic effects on pupal phenotypes, were not picked up by GWA. Actually, there is no observed significantly lower genetic density or allele frequency of random chosen genes with significant pupal case length phenotypic effects compared with GWA candidate genes (Figure 6-figure supplement 2). Therefore, the reason why they have not shown up in the GWA has to be that they do not include segregating variants of sufficient effect size in the population from which the DGRP was derived. In fact, one can expect that most genetic variants present in a natural *Drosophila* population are unlikely to represent gene disabling mutations. And the variants that are gene disabling should be rare because they should be selected against, *i.e*., they should also only seldomly occur as homozygotes. Hence, to test these genes as homozygous gene disruptions is rather unnatural. Still, it is an indication that these genes are involved in some form in the phenotype. This is a general issue when comparing gene effects from a GWA analysis with those from classic genetic analyses. The former trace the effects of naturally occurring variants, the latter the ones of gene disruptions. These converge only for human genetic disease studies, but not for studies on natural genetic variation of quantitative traits. One has to keep this dichotomy of genetic views in mind when placing GWA results in the context of classic genetic results.

## Conclusion

Our data confirm the major prediction of the polygenic/omnigenic model, namely that most randomly chosen genes can potentially influence any given trait. It should be possible to apply this test also to other phenotypes or genetic systems where the necessary stocks or experimental procedures are available. Evidently, if almost any random gene is involved in a given phenotype, why should one then do a GWA in the first place? Hence, it should be of special interest to assess whether GWA p-values provide generally a guide towards the core networks required to generate the phenotype.

## Materials and Methods

### Drosophila stocks and flies rearing

All flies were reared in vials containing cornmeal–molasses–agar medium at 24°C, 55% -78% relative humidity, and a 12:12-h light-dark cycle. A HOBO® data logger was placed in the incubator to monitor and record any potential environmental changes, including temperature, light and humidity. Given the only minor impact on pupal case length from the change of relative humidity across experiments (Figure 2-figure supplement 1B), no special action was taken to control this factor.

In total, 14 wild type stocks, 198 DGRP inbred lines, 58 transposon insertion mutagenesis stocks (disruption for 55 annotated protein coding genes) and 3 corresponding progenitor stocks were assayed in this study. The detailed description and measurement data of these stocks can be found in Supplementary Table S1.

### Phenotyping of pupal case length

We conducted the measurement of pupal case length by adopting an earlier established image-analysis based automated phenotyping pipeline as described in (Reeves and Tautz, 2017; Zhang et al., 2020). The entire phenotyping pipeline is implemented with five successive procedures:

#### 1) Vial preparation

Standard cornmeal–molasses–agar food was dispensed into standard 28.5 mm diameter, 95 mm height vials (Genesee Scientific) to a depth of approximately 20mm. Once the food vials had fully cooled, 10.1 cm x 10.5 cm squares of transparent film (nobo, plain paper copier film, 33638237) were slid into the bottom of each vial lining their entire vertical wall. Inserting the film does not change the storage properties of the food. Prior to introducing adult flies, a very small amount of live yeast paste was dotted on the food surface. A barcode with custom printed semi-transparent label was affixed to the outside of each vial as the unique identifier.

#### 2) Mating flies

Approximately ten 2-5 days old healthy female flies (fifteen for inbred stocks to compensate the reduced fertility) and five similar-aged male flies were introduced into each vial, under the incubation condition as mentioned above. These adult flies were cleared from the vials after 1-2 days introduction and vials were kept in the same incubation condition for another 8-9 days to allow them to reach pupation stage. In case of observing that most of offspring in the vials were present as pupae attached to the transparent film, the film was then gently taken out from each vial, and the food from the lower part was scrapped away and any viable larvae were also removed.

#### 3) Film Photographing

Once removed from the vial, the film (with the barcode label affixed) was then placed into a custom plastic frame, which holds the film flat for further photographing. Frames were then photographed in a light tight box with a sliding door, which provided illumination only from underneath the frame, effectively silhouetting the pupae while minimizing tangential shadows. Every photograph included a 1 euro cent coin (16.25 mm diameter), for the control of camera coordinate changes and the conversion of measurements from pixels to millimetres (mm). Batches of the resulting images were then introduced into the following image analysis procedure.

#### 4) Image analysis

We applied the open-source image analysis software CellProfiler (v2.1.0) (Carpenter et al., 2006) for the recognition of pupae and measurements of a variety of attributes, with a customized pipeline adopted from (Zhang et al., 2020). Firstly, by using the CellProfiler module “identify primary objects”, we identified any “primary object” with significant distinction from the background without restriction on their sizes. Secondly, the above identified objects composing of multiple touching pupae were disentangled into distinct pupae (module “Untangle Worms”). Thirdly, the resulting candidate pupae were shrunk and re-propagated outwards for a more precise detection of the edges of each pupa based on boundary changes in pixel intensity (module “Identify secondary objects”). Lastly, distinct attributes for the pupae were calculated and a specific confidence class was assigned for each pupa based on its size attribute. Based on manual curation on 40 randomly selected films, we found that the above CellProfiler pipeline can reach to a 96% of sensitivity (fraction of identified true pupae), but a modest false discovery rate of 19% for identified putative pupae (Zhang et al., 2020). To further reduce the false discovery rate, we refined an additional criterion setting based on the size attributes of “true” pupae based on manual curation (Zhang et al., 2020). Applying this setting of new criteria, the false discovery rate for pupae detection dropped to around 0.15%, with only a tiny fraction (< 0.7%) of loss for true positive results.

#### 5) Pupal case length measurement

The pupal case length is defined as the length of the major axis of the ellipse that has the same normalized second central moments as the region of identified pupae, measured with the “Areashape_MajorAxisLength” index in CellProfiler. Based on its ratio to the diameter measurement of 1 euro cent coin (16.25 mm diameter) included in the photograph, the measurement of case length of each pupa was converted from pixels to mm. The mean of all the measurements of pupal case length for all the pupae in the food vial was taken as representative for the focal vial. To further reduce the potential bias from low sampling effect, we only included vials with a pupal density of a minimum of 15, and required each stock should include at least 6 such reliable vial measurements.

### Repeated measurements of control stocks

Throughout the experiments for wildtype and DGRP inbred strains measurements, two wild-type stocks (S-314 and S-317) were continually re-measured in the manner described above to control for environmental effects, especially small fluctuations in humidity. Such re-measurement of control stocks were not applied for the functional validation experiments as shown below, as the experimental tests on each pair of gene disruption stock and progenitor stock were conducted in the same experimental condition. The impacts of the incubator relative humidity changes on the pupal case length measurement (across all rounds of DGRP inbred stock experiments) were directly examined by using Pearson’s correlation tests for two control stocks separately.

### Wolbachia infection effect test

*Wolbachia pipientis* is a maternally transmitted endosymbiotic bacterium that was reported to infect around 53% of DGRP inbred stocks (Supplementary Table S1b) (Huang et al., 2014). We used two different approaches to examine whether there is any significant influence on pupal case length from *Wolbachia* infection. Firstly, we directly compared the pupal case length measurement between *Wolbachia*-infected stocks and *Wolbachia*-uninfected stocks, and computed the statistical significance on the pupal case length difference between these two groups via a Wilcoxon rank-sum test.

Secondly, we conducted a direct experimental test to compare the changes of pupal case length measurement between *Wolbachia* infected stocks and *Wolbachia*-free stocks, which were created though two generations of tetracycline treatment on the infected stocks (by adding an appropriate volume of 100 mg/ml of tetracycline suspended in 99% ethanol to the surface of the solid prepared food) and then reared for at least another two generations with standard food to avoid any detrimental parental effects (Zeh et al., 2012). We firstly ruled out the possibility that tetracycline treatment could have an influence on pupal case length, by comparing the pupal case length measurement changes between three randomly selected *Wolbachia*-free stocks (DR_14, DR_45, DR_106), and the same stocks with the above mentioned tetracycline treatment procedure. Then we randomly selected three *Wolbachia* infected stocks (DR_16, DR_21, DR_67), and half of the files were treated with tetracycline as mentioned to create *Wolbachia*-free stocks, and the rest half the flies from same three strains were also reared with standard food across the experiment as controls. We extracted genomic DNA from the above 6 stocks individually by using DNeasy blood and tissue kit (Qiagen), and measured the purity and concentration of the resulting DNA with NanoDrop ND-1000 spectrophotometer (Thermofisher). A diagnostic PCR to test for the presence of the *Wolbachia wsp* gene was performed by using the primers wsp81F (5’-tggtccaaaatgtgagaaac-3’) and wsp691r (5’-aaaattaaacgctactcca-3’) (Richardson et al., 2012), under the reaction condition of 35 cycles of 94°C for 15 seconds, 55°C for 30 seconds and 72°C for 1 minute. A standard (1%) agarose gel electrophoresis was used to test for the presence of the PCR product (∼630 bp), with the broad range Quick DNA Marker (NEB #N0303) as loading ladder. Pupal case length measurement between the three *Wolbachia*-infected and created *Wolbachia*-free lines were then measured and compared by using Wilcoxon rank-sum tests.

### Principal component analysis and genomic inversion effect test

All the genetic variant calling data and major genomic inversion status were retrieved from DGRP freeze 2 (Huang et al., 2014), for which the coordinates were based on Flybase version 5 (Attrill et al., 2016). Only the genetic variants with <20% missing values and ≥5% minor allele frequency (MAF) were kept (corresponding to 1,903,028 genetic variants) for further analysis. We assessed the possible influence of population structure on the pupal case length measurement in DGRP inbred lines, by using the PCA module from PLINK v1.90 (Purcell et al., 2007) to identify principal components (PCs) from the above filtered genetic variant data. We firstly checked the clustering status on the three equal-sized groups of DGRP inbred strains with ranked pupal case length values (*i.e.*, low pupal case length value, medium pupal case length value and high pupal case length value) based on the projection analysis of the top two PCs, and then examined the correlation between pupal case length values and the top 20 PCs by using Pearson’s correlation tests.

To explore the potential contribution of major genomic inversions to the population structure within DGRP strains, we used Pearson’s correlation tests to calculate the correlations between genomic inversion status and the top 2 PCs from the above principal component analysis. Moreover, we also tested the correlation between pupal case length values and the status of each genomic inversion within DGRP strains. The resultant significant correlated major genomic inversion (*i.e.*, In(3R)Mo) was included as a covariate for the following chromosomal effect partitioning and genome-wide association analysis.

### Estimates of heritability and chromosomal effect partitioning

We estimated the broad sense heritability (H^2^) of pupal case length with the variance components of a linear model of the form: Phenotype = Population mean + Line effect + error (Schmidt et al., 2017). We computed the total phenotypic variance as Genetic Variance (G_v_) + Environmental Variance (E_v_), and the H^2^ as G_v_/G_v_+E_v_. This calculations for both wild-type and DGRP inbred stocks were implemented by using IBM SPSS Statistics (version 22) (SPSS Inc, 2007), with pupal case length measurement as the dependent variable and DGRP stock names as a random factor. Meanwhile, we compared the estimates of H^2^ computed with the same methodology, but from different *Drosophila melanogaster* strains datasets (Reeves and Tautz, 2017). The estimates on the narrow sense heritability (h^2^) of pupal case length based on mid-parent regression were directly retrieved from (Reeves and Tautz, 2017).

We partitioned the fraction of pupal case length phenotypic variance explained by the genetic variants from each *Drosophila melanogaster* chromosome/arm (*i.e.*, 2L, 2R, 3L, 3R, 4, X) by using GEMMA (version 0.96) (Zhou and Stephens, 2012). GEMMA implemented a univariate linear mixed model to estimate the variance (denoted as PVE) explained by each chromosome/arm, with the controlling for population structure effect via incorporating a genetic relationship matrix (GRM) built from DGRP strains and the major genomic inversion In(3R)Mo status as a covariate in the analysis. Even though the absolute estimates are inflated due to the individual relatedness, the relative effects of chromosomes/arms are still informative for comparison, as the overestimation is uniformly spread across the whole genome (Pallares et al., 2014; Yang et al., 2011).

### Genome-wide association analysis

We performed genome-wide association (GWA) analysis on pupal case length for the above filtered genetic variants from DGRP freeze 2 (Huang et al., 2014), by using the FastLMM (Lippert et al., 2011) program (version 0.2.32), with the major genomic inversion In(3R)Mo status being included as a covariate for the GWA analysis. This program fitted a linear mixed model that could control for population stratification effects.

We defined GWA significant associated genetic variants by using p-value < 1 × 10^−5^, which is a nominal threshold frequently used in *Drosophila* quantitative trait genetic studies (Dembeck et al., 2015; Lee et al., 2017; Zhang et al., 2020). The R package “qqman” (Turner, 2014) was exploited for the visualization of GWA results in a Manhattan plot and QQ plot. We calculated the effect size of each significant genetic variant in the following way (Pallares et al., 2014):

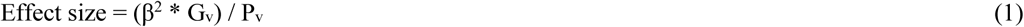

Where G_v_ is the variance of the genotype at the focal genetic variant, and P_v_ is the total variance of the pupal case length phenotype. β is reported as “SNPWeight” for each genetic variant in the FastLMM output.

We predicted the effects of GWA significant genetic variants by using SnpEff (Cingolani et al., 2012) with default parameters, and taken all the corresponding affected protein-coding genes as associating genes. In short, all the protein-coding genes within 5 kb up/down-stream of focal genetic variant were taken as its associating genes. We also tested the genotypic linkage disequilibrium (LD) for each GWA significant genetic variant and other genetic variants by calculating the squared correlation estimator r^2^ with PLINK v1.90 (Purcell et al., 2007). A significant genetic region (QTL) was defined by the position of the most distant downstream and upstream genetic variants showing a minimum r^2^ of 0.8 to the focal GWA significant genetic variant. The combined associating genes from SnpEff predictions and the genes within the QTL regions were considered as GWA candidate genes.

### Expression analysis

We downloaded the *Drosophila melanogaster* developmental raw Illumina paired-end RNA-Seq data from Graveley et al (Graveley et al., 2011) (NCBI SRA accession: SRP001065). This dataset included the transcriptome of 27 distinct developmental stages covering all the major steps in *Drosophila* life cycle, which were further collapsed into six super-developmental stages: *1) Early embryo*: including 0-2 hrs. embryo, 2-4 hrs. embryo, 4-6 hrs. embryo, and 6-8 hrs. embryo; *2) Middle embryo:* including 8-10 hrs. embryo, 10-12 hrs. embryo, 12-14 hrs. embryo, and 14-16 hrs. embryo; *3) Late embryo:* including 16-18 hrs. embryo, 18-20 hrs. embryo, 20-22 hrs. embryo, and 22-24 hrs. embryo; *4) Larval stage:* including L1 stage, L2 stage, L3 stage of 12 hr. post-molt, L3 stage of dark blue gut (PS1-2), L3 stage of light blue gut (PS3-6), and L3 stage of clear blue gut (PS7-9); *5) Pupal stage:* including white prepupa (WPP), 12 hr. after WPP (P5), 24 hr. after WPP (P6), 2 days after WPP (P8), 3 days after WPP (P9-10), and 4 days after WPP (P15); *6) Adult stage:* male/female after 1-day eclosion, male/female after 5-day eclosion, and male/female after 30-day eclosion.

We trimmed and filtered the low-quality raw Illumina RNA-Seq reads sequenced from the sample of each developmental stage by using Fastp program v0.20.0 (Chen et al., 2018), and only included the sequencing reads with minimum length of 20bp and average quality score of 20 for further analysis. The filtered RNA-Seq reads were aligned to *Drosophila melanogaster* v6 reference genome sequence with HISAT2 v2.1.0 (Kim et al., 2019), taking advantage of the protein-coding gene annotation in Flybase v6.32 (Attrill et al., 2016) by using the --exon option of the hisat2-build. Then we counted the fragments mapped to the annotated genes with featureCounts v1.6.3 (Liao et al., 2014), and calculated the expression level of each annotated coding gene from the sample of each developmental stage in the unit of FPKM (Fragments Per Kilobase of transcript per Million mapped reads).

We calculated the gene expression enrichment fold changes across these 27 developmental stages as the ratios between the fractions of genes with expression (FPKM >0) for GWA candidate genes and that of total protein coding gene annotated in Flybase v6.32 (Attrill et al., 2016). We took the average expression enrichment statistics of males and females in the adult stages as for representation. The statistical significances (p-values) of expression enrichment patterns were computed by using Fisher’s exact tests.

### Automatic measurement of pupation site choice

We also assessed the pupation height (an indicator of pupation site choice) for the gene disruption stocks used in the experimental validation tests as shown below. The measurement of pupation height follows the procedure described in (Zhang et al., 2020). In brief, the pupation height was defined as the distance from the vertical coordinate of pupation site (pupal centre) to the food surface in the vial in millimetre (mm). As the pupation height measurements are sensitive to the pupal density in the vial, the raw measurements for pupation height were further adjusted according to the equations described in (Zhang et al., 2020).

### Functional validation experiments

We validated the phenotypic effects of 10 GWA candidate genes from 11 gene disruption mutagenesis stocks (The gene *Drl-2* was tested twice with different disruption locations), based on their stock availability in the Drosophila gene disruption project (Bellen et al., 2011) (Supplementary Table S1c). All these mutagenesis stocks were conducted with transposon insertion disruption, for which each focal gene was disrupted by a transposon (Minos, P-element or PBac) insertion in the gene region (either coding or regulatory region). The exact landing site of the transposon was determined based on the sequencing reads of both sides of flanking regions (Bellen et al., 2011). The detailed information about these gene mutagenesis stocks and their co-isogenic progenitors is provided in Supplementary Table S1c. Except one disruption stock (Blooming stock ID: 19060; disrupted gene: *Pxn*), all the rest 10 stocks are homozygous insertion viable, and thus were tested as homozygotes (In case of semi-lethal, only the adult flies with no balancer maker were used for experimental tests). For each test, at least 8 vials (15 females and 5 males introduced to each vial) were set up for each insertion and its progenitor stock, and their phenotypic values (pupal case length and pupation height) were measured by following the above phenotyping pipeline. The experimental tests on each pair of gene disruption stock and progenitor stock were conducted in the same experimental round, thus any potential influence (food preparation, incubator environment, et al) on the phenotype measurements can be avoided. The experimental effect size of each gene disruption was calculated as the average deviation of phenotypic values of gene disruption stock from that of its progenitor stock. The statistical differences of the phenotypic measurements between gene disruption stocks and their progenitor stocks were performed by using Wilcoxon rank-sum tests.

We developed a distinct approach to test the phenotypic effect of the above homozygous complete-lethal stock (Zhang et al., 2020), which is segregating balancer chromosomes with Tubby (Tb) as visible marker at pupal case (short rounded pupae). We crossed this gene disruption stock (virgin females) with its progenitor stock (males) to generate F1 generation, and only measured the phenotypic values of hemizygous insertion individuals without Tb maker (no presence of balancer chromosome), and then compared with that of its progenitor stock. The presence/absence of Tb maker for individual pupae was distinguished by their pupal case length (>73 pixels for Areashape_major_len from the output of CellProfiler), on the observation of the clear distinction between individual pupae with and without Tb marker (Zhang et al., 2020). Moreover, a manual curation was performed to further separate ambiguous individuals. It should be noted that the phenotypic effect of this homozygous complete-lethal stock was tested in hemizygote manner (Blooming stock ID: 19060; disrupted gene: *Pxn*).

Additionally, we also experimentally tested the phenotypic effects of 45 selected genes, on the basis of the following criteria: 1) the GWA p-value on the focal gene falls out the significance threshold (1 × 10^−5^); 2) the focal gene has detectable expression in relevant development stages for pupal case length (late larval and pupal stage; FPKM >0), and 3) the focal gene has homozygous transposon insertion disruption viable stock(s) from the panel of *Drosophila* gene disruption project (Bellen et al., 2011). The latter criterion biases against essential genes (approximately 30% are not homozygous viable), but otherwise the selection was essentially random. Two of the above chosen genes (*CG14007*, and *CG42260*) have 2 qualified transposon insertion stocks with disruption in different locations (while one disruption stock for each of the rest 43 genes), and thus in total 47 transposon insertion stocks were applied for the experimental tests. The phenotyping test experiments were conducted with the same procedure as stated above. The GWA association p-value was taken as the lowest p-values from all the genetic variants within the gene and the 5kb up/down-stream of target gene.

Furthermore, we analysed the pattern on absolute pupal case length phenotypic effect size (absolute deviation of the phenotypic values of gene disruption stock from that of progenitor stock) of the disruption of the genes, based on their categories: 1) GWA candidate gene or randomly chosen gene; 2) the gene region of the transposon insertion landing site (coding region, intron or UTR); 3) the percentage of disruption on gene’s coding region. For the latter one, only the gene disruptions in the coding region were included for analysis, and the average percentage of disruption was taken as representative, in case of multiple transcripts for a focal gene. The statistical difference on the absolute pupal case length phenotypic effect size between two categorized groups were tested by using Wilcoxon rank-sum tests, and the correlation between the absolute pupal case length phenotypic effect size and percentage of gene’s coding region disruption with Pearson’s correlation test.

### Genetic variant density analysis

We compared the genetic variant density (# of genetic variants per kb region) pattern between GWA candidate genes and randomly chosen genes (the subset which have significant phenotypic effects on pupal case length, p-value⩽0.05), based on the information from of DGRP freeze 2 (Huang et al., 2014). The genetic variants within 5kp up/downstream the focal gene (include the gene itself) were included for the analysis. Two types of genetic variants were analysed: 1) the total genetic variants; 2) the “good” genetic variants that passed the criteria for GWA analysis, *i.e.*, with missing values below 20% and minor allele frequency above 5%. For the latter one, we also compared the minor allele frequency (MAF) pattern between these two groups of genes. Overall comparisons of genetic variant density between two groups of genes (whether GWA candidate genes have on average higher variant density/MAF than random genes) were performed with one-sided Wilcoxon rank-sum test.

### Data availability

All the *Drosophila melanogaster* strains used in this study are public available through either Bloomington *Drosophila* Stock centre (https://bdsc.indiana.edu/) or EHIME *Drosophila* stock centre (https://kyotofly.kit.jp/cgi-bin/ehime/index.cgi). The detailed information about the global wildtype, DGRP inbred lines, gene disruption stocks and their progenitors can be found in the Supplementary Table 1. The primers used for *Wolbachia* infection detection are listed in the above text of this section.

## Acknowledgements

We thank the lab members for helpful discussions and suggestions. We thank Anita Moeller, Elke Blohm Sievers, and Michaela Schwarz for their excellent technical help with conducting this experiment. This work was supported by institutional funding through the Max-Planck Society.

## Supporting information

**Supplementary Table 1:** Information for the fly stocks used in this study

## References

Attrill H, Falls K, Goodman JL, Millburn GH, Antonazzo G, Rey AJ, Marygold SJ. 2016. Flybase: Establishing a gene group resource for Drosophila melanogaster. Nucleic Acids Res 44:D786–792. doi: 10.1093/nar/gkv1046

Bellen HJ, Levis RW, He Y, Carlson JW, Evans-Holm M, Bae E, Kim J, Metaxakis A, Savakis C, Schulze KL, Hoskins RA, Spradling AC. 2011. The Drosophila gene disruption project: Progress using transposons with distinctive site specificities. Genetics 188:731–743. doi: 10.1534/genetics.111.126995

Boyle EA, Li YI, Pritchard JK. 2017. An Expanded View of Complex Traits: From Polygenic to Omnigenic. Cell 169:1177–1186. doi: 10.1016/j.cell.2017.05.038

Carpenter AE, Jones TR, Lamprecht MR, Clarke C, Kang IH, Friman O, Guertin DA, Chang JH, Lindquist RA, Moffat J, Golland P, Sabatini DM. 2006. CellProfiler: Image analysis software for identifying and quantifying cell phenotypes. Genome Biol 7:R100. doi: 10.1186/gb-2006-7-10-r100

Chen S, Zhou Y, Chen Y, Gu J. 2018. Fastp: An ultra-fast all-in-one FASTQ preprocessor. Bioinformatics 34:i884–i890. doi: 10.1093/bioinformatics/bty560

Cingolani P, Platts A, Wang LL, Coon M, Nguyen T, Wang L, Land SJ, Lu X, Ruden DM. 2012. A program for annotating and predicting the effects of single nucleotide polymorphisms, SnpEff: SNPs in the genome of Drosophila melanogaster strain w1118; iso-2; iso-3. Fly (Austin) 6:80–92. doi: 10.4161/fly.19695

Dembeck LM, Böröczky K, Huang W, Schal C, Anholt RR, Mackay TF. 2015. Genetic architecture of natural variation in cuticularhydrocarbon composition in Drosophila melanogaster. Elife 4:e09861. doi: 10.7554/eLife.09861

Durham MF, Magwire MM, Stone EA, Leips J. 2014. Genome-wide analysis in Drosophila reveals age-specific effects of SNPs on fitness traits. Nat Commun 5:4338. doi: 10.1038/ncomms5338

Graveley BR, Brooks AN, Carlson JW, Duff MO, Landolin JM, Yang L, Artieri CG, Van Baren MJ, Boley N, Booth BW, Brown JB, Cherbas L, Davis CA, Dobin A, Li R, Lin W, Malone JH, Mattiuzzo NR, Miller D, Sturgill D, Tuch BB, Zaleski C, Zhang D, Blanchette M, Dudoit S, Eads B, Green RE, Hammonds A, Jiang L, Kapranov P, Langton L, Perrimon N, Sandler JE, Wan KH, Willingham A, Zhang Y, Zou Y, Andrews J, Bickel PJ, Brenner SE, Brent MR, Cherbas P, Gingeras TR, Hoskins RA, Kaufman TC, Oliver B, Celniker SE. 2011. The developmental transcriptome of Drosophila melanogaster. Nature 471:473–479. doi: 10.1038/nature09715

Hoffmann AA, Rieseberg LH. 2008. Revisiting the Impact of Inversions in Evolution: From Population Genetic Markers to Drivers of Adaptive Shifts and Speciation? Annu Rev Ecol Evol Syst 39:21–42. doi: 10.1146/annurev.ecolsys.39.110707.173532

Huang W, Massouras A, Inoue Y, Peiffer J, Ràmia M, Tarone AM, Turlapati L, Zichner T, Zhu D, Lyman RF, Magwire MM, Blankenburg K, Carbone MA, Chang K, Ellis LL, Fernandez S, Han Y, Highnam G, Hjelmen CE, Jack JR, Javaid M, Jayaseelan J, Kalra D, Lee S, Lewis L, Munidasa M, Ongeri F, Patel S, Perales L, Perez A, Pu LL, Rollmann SM, Ruth R, Saada N, Warner C, Williams A, Wu YQ, Yamamoto A, Zhang Y, Zhu Y, Anholt RRH, Korbel JO, Mittelman D, Muzny DM, Gibbs RA, Barbadilla A, Johnston JS, Stone EA, Richards S, Deplancke B, Mackay TFC. 2014. Natural variation in genome architecture among 205 Drosophila melanogaster Genetic Reference Panel lines. Genome Res 24:1193–1208. doi: 10.1101/gr.171546.113

Jones MDR, Reiter P. 1975. Entrainment of the pupation and adult activity rhythms during development in the mosquito Anopheles gambiae. Nature 254:242–244. doi: 10.1038/254242a0

Kim D, Paggi JM, Park C, Bennett C, Salzberg SL. 2019. Graph-based genome alignment and genotyping with HISAT2 and HISAT-genotype. Nat Biotechnol 37:907–915. doi: 10.1038/s41587-019-0201-4

Lattao R, Bonaccorsi S, Guan X, Wasserman SA, Gatti M. 2011. Tubby-tagged balancers for the drosophila X and second chromosomes. Fly (Austin) 5:369–370. doi: 10.4161/fly.5.4.17283

Lee YCG, Yang Q, Chi W, Turkson SA, D. WA, Kemkemer C, Zeng ZB, Long M, Zhuang X. 2017. Genetic architecture of natural variation underlying adult foraging behavior that is essential for survival of Drosophila melanogaster. Genome Biol Evol 9:1357–1369. doi: 10.1093/gbe/evx089

Liao Y, Smyth GK, Shi W. 2014. FeatureCounts: An efficient general purpose program for assigning sequence reads to genomic features. Bioinformatics 30:923–930. doi: 10.1093/bioinformatics/btt656

Lippert C, Listgarten J, Liu Y, Kadie CM, Davidson RI, Heckerman D. 2011. FaST linear mixed models for genome-wide association studies. Nat Methods 8:833–835. doi: 10.1038/nmeth.1681

Liu X, Li YI, Pritchard JK. 2019. Trans Effects on Gene Expression Can Drive Omnigenic Inheritance. Cell 177:1022–1034. doi: 10.1016/j.cell.2019.04.014

Loubéry S, Seum C, Moraleda A, Daeden A, Fürthauer M, Gonzalez-Gaitan M. 2014. Uninflatable and notch control the targeting of sara endosomes during asymmetric division. Curr Biol 25:817–818. doi: 10.1016/j.cub.2014.07.054

Mackay TFC, Huang W. 2018. Charting the genotype–phenotype map: lessons from the Drosophila melanogaster Genetic Reference Panel. Wiley Interdiscip Rev Dev Biol 7:e289. doi: 10.1002/wdev.289

MacKay TFC, Richards S, Stone EA, Barbadilla A, Ayroles JF, Zhu D, Casillas S, Han Y, Magwire MM, Cridland JM, Richardson MF, Anholt RRH, Barrón M, Bess C, Blankenburg KP, Carbone MA, Castellano D, Chaboub L, Duncan L, Harris Z, Javaid M, Jayaseelan JC, Jhangiani SN, Jordan KW, Lara F, Lawrence F, Lee SL, Librado P, Linheiro RS, Lyman RF, MacKey AJ, Munidasa M, Muzny DM, Nazareth L, Newsham I, Perales L, Pu LL, Qu C, Ràmia M, Reid JG, Rollmann SM, Rozas J, Saada N, Turlapati L, Worley KC, Wu YQ, Yamamoto A, Zhu Y, Bergman CM, Thornton KR, Mittelman D, Gibbs RA. 2012. The Drosophila melanogaster Genetic Reference Panel. Nature 482:173–178. doi: 10.1038/nature10811.

Pallares LF, Harr B, Turner LM, Tautz D. 2014. Use of a natural hybrid zone for genomewide association mapping of craniofacial traits in the house mouse. Mol Ecol 23:5756–5770. doi: 10.1111/mec.12968

Price GM. 1970. Pupation Inhibiting Factor in the Larva of the Blowfly Calliphora erythrocephala. Nature 228:876–877. doi: 10.1038/228876a0

Purcell S, Neale B, Todd-Brown K, Thomas L, Ferreira MAR, Bender D, Maller J, Sklar P, de Bakker PIW, Daly MJ, Sham PC. 2007. PLINK: A Tool Set for Whole-Genome Association and Population-Based Linkage Analyses. Am J Hum Genet 81:559–575. doi: 10.1086/519795

Reeves RG, Tautz D. 2017. Automated Phenotyping Indicates Pupal Size in Drosophila Is a Highly Heritable Trait with an Apparent Polygenic Basis. G3 (Bethesda) 7:1277–1286. doi: 10.1534/g3.117.039883

Richardson MF, Weinert LA, Welch JJ, Linheiro RS, Magwire MM, Jiggins FM, Bergman CM. 2012. Population Genomics of the Wolbachia Endosymbiont in Drosophila melanogaster. PLoS Genet 8:e1003129. doi: 10.1371/journal.pgen.1003129

Sakurai M, Aoki T, Yoshikawa S, Santschi LA, Saito H, Endo K, Ishikawa K, Kimura KI, Ito K, Thomas JB, Hama C. 2009. Differentially expressed Drl and Drl-2 play opposing roles in Wnt5 signaling during drosophila olfactory system development. J Neurosci 29:4972–4980. doi: 10.1523/JNEUROSCI.2821-08.2009

Schmidt JM, Battlay P, Gledhill-Smith RS, Good RT, Lumb C, Fournier-Level A, Robin C. 2017. Insights into DDT resistance from the drosophila melanogaster genetic reference panel. Genetics 207:1181–1193. doi: 10.1534/genetics.117.300310

SPSS Inc. 2007. SPSS Advanced Statistics 17.0. Statistics (Ber).

Turner SD. 2014. qqman: an R package for visualizing GWAS results using Q-Q and manhattan plots, bioRxiv. doi: 10.1101/005165

Wood AR, Esko T, Yang J, Vedantam S, Pers TH, Gustafsson S, Chu AY, Estrada K, Luan J, Kutalik Z, Amin N, Buchkovich ML, Croteau-Chonka DC, Day FR, Duan Y, Fall T, Fehrmann R, Ferreira T, Jackson AU, Karjalainen J, Lo KS, Locke AE, Mägi R, Mihailov E, Porcu E, Randall JC, Scherag A, Vinkhuyzen AAE, Westra HJ, Winkler TW, Workalemahu T, Zhao JH, Absher D, Albrecht E, Anderson D, Baron J, Beekman M, Demirkan A, Ehret GB, Feenstra B, Feitosa MF, Fischer K, Fraser RM, Goel A, Gong J, Justice AE, Kanoni S, Kleber ME, Kristiansson K, Lim U, Lotay V, Lui JC, Mangino M, Leach IM, Medina-Gomez C, Nalls MA, Nyholt DR, Palmer CD, Pasko D, Pechlivanis S, Prokopenko I, Ried JS, Ripke S, Shungin D, Stancáková A, Strawbridge RJ, Sung YJ, Tanaka T, Teumer A, Trompet S, Van Der Laan SW, Van Setten J, Van Vliet-Ostaptchouk J V., Wang Z, Yengo L, Zhang W, Afzal U, Ärnlöv J, Arscott GM, Bandinelli S, Barrett A, Bellis C, Bennett AJ, Berne C, Blüher M, Bolton JL, Böttcher Y, Boyd HA, Bruinenberg M, Buckley BM, Buyske S, Caspersen IH, Chines PS, Clarke R, Claudi-Boehm S, Cooper M, Daw EW, De Jong PA, Deelen J, Delgado G, Denny JC, Dhonukshe-Rutten R, Dimitriou M, Doney ASF, Dörr M, Eklund N, Eury E, Folkersen L, Garcia ME, Geller F, Giedraitis V, Go AS, Grallert H, Grammer TB, Gräßler J, Grönberg H, De Groot LCPGM, Groves CJ, Haessler J, Hall P, Haller T, Hallmans G, Hannemann A, Hartman CA, Hassinen M, Hayward C, Heard-Costa NL, Helmer Q, Hemani G, Henders AK, Hillege HL, Hlatky MA, Hoffmann W, Hoffmann P, Holmen O, Houwing-Duistermaat JJ, Illig T, Isaacs A, James AL, Jeff J, Johansen B, Johansson Å, Jolley J, Juliusdottir T, Junttila J, Kho AN, Kinnunen L, Klopp N, Kocher T, Kratzer W, Lichtner P, Lind L, Lindström J, Lobbens S, Lorentzon M, Lu Y, Lyssenko V, Magnusson PKE, Mahajan A, Maillard M, McArdle WL, McKenzie CA, McLachlan S, McLaren PJ, Menni C, Merger S, Milani L, Moayyeri A, Monda KL, Morken MA, Müller G, Müller-Nurasyid M, Musk AW, Narisu N, Nauck M, Nolte IM, Nöthen MM, Oozageer L, Pilz S, Rayner NW, Renstrom F, Robertson NR, Rose LM, Roussel R, Sanna S, Scharnagl H, Scholtens S, Schumacher FR, Schunkert H, Scott RA, Sehmi J, Seufferlein T, Shi J, Silventoinen K, Smit JH, Smith AV, Smolonska J, Stanton A V., Stirrups K, Stott DJ, Stringham HM, Sundström J, Swertz MA, Syvänen AC, Tayo BO, Thorleifsson G, Tyrer JP, Van Dijk S, Van Schoor NM, Van Der Velde N, Van Heemst D, Van Oort FVA, Vermeulen SH, Verweij N, Vonk JM, Waite LL, Waldenberger M, Wennauer R, Wilkens LR, Willenborg C, Wilsgaard T, Wojczynski MK, Wong A, Wright AF, Zhang Q, Arveiler D, Bakker SJL, Beilby J, Bergman RN, Bergmann S, Biffar R, Blangero J, Boomsma DI, Bornstein SR, Bovet P, Brambilla P, Brown MJ, Campbell H, Caulfield MJ, Chakravarti A, Collins R, Collins FS, Crawford DC, Cupples LA, Danesh J, De Faire U, Den Ruijter HM, Erbel R, Erdmann J, Eriksson JG, Farrall M, Ferrannini E, Ferrières J, Ford I, Forouhi NG, Forrester T, Gansevoort RT, Gejman P V., Gieger C, Golay A, Gottesman O, Gudnason V, Gyllensten U, Haas DW, Hall AS, Harris TB, Hattersley AT, Heath AC, Hengstenberg C, Hicks AA, Hindorff LA, Hingorani AD, Hofman A, Hovingh GK, Humphries SE, Hunt SC, Hypponen E, Jacobs KB, Jarvelin MR, Jousilahti P, Jula AM, Kaprio J, Kastelein JJP, Kayser M, Kee F, Keinanen-Kiukaanniemi SM, Kiemeney LA, Kooner JS, Kooperberg C, Koskinen S, Kovacs P, Kraja AT, Kumari M, Kuusisto J, Lakka TA, Langenberg C, Le Marchand L, Lehtimäki T, Lupoli S, Madden PAF, Männistö S, Manunta P, Marette A, Matise TC, McKnight B, Meitinger T, Moll FL, Montgomery GW, Morris AD, Morris AP, Murray JC, Nelis M, Ohlsson C, Oldehinkel AJ, Ong KK, Ouwehand WH, Pasterkamp G, Peters A, Pramstaller PP, Price JF, Qi L, Raitakari OT, Rankinen T, Rao DC, Rice TK, Ritchie M, Rudan I, Salomaa V, Samani NJ, Saramies J, Sarzynski MA, Schwarz PEH, Sebert S, Sever P, Shuldiner AR, Sinisalo J, Steinthorsdottir V, Stolk RP, Tardif JC, Tönjes A, Tremblay A, Tremoli E, Virtamo J, Vohl MC, Amouyel P, Asselbergs FW, Assimes TL, Bochud M, Boehm BO, Boerwinkle E, Bottinger EP, Bouchard C, Cauchi S, Chambers JC, Chanock SJ, Cooper RS, De Bakker PIW, Dedoussis G, Ferrucci L, Franks PW, Froguel P, Groop LC, Haiman CA, Hamsten A, Hayes MG, Hui J, Hunter DJ, Hveem K, Jukema JW, Kaplan RC, Kivimaki M, Kuh D, Laakso M, Liu Y, Martin NG, März W, Melbye M, Moebus S, Munroe PB, Njølstad I, Oostra BA, Palmer CNA, Pedersen NL, Perola M, Pérusse L, Peters U, Powell JE, Power C, Quertermous T, Rauramaa R, Reinmaa E, Ridker PM, Rivadeneira F, Rotter JI, Saaristo TE, Saleheen D, Schlessinger D, Slagboom PE, Snieder H, Spector TD, Strauch K, Stumvoll M, Tuomilehto J, Uusitupa M, Van Der Harst P, Völzke H, Walker M, Wareham NJ, Watkins H, Wichmann HE, Wilson JF, Zanen P, Deloukas P, Heid IM, Lindgren CM, Mohlke KL, Speliotes EK, Thorsteinsdottir U, Barroso I, Fox CS, North KE, Strachan DP, Beckmann JS, Berndt SI, Boehnke M, Borecki IB, McCarthy MI, Metspalu A, Stefansson K, Uitterlinden AG, Van Duijn CM, Franke L, Willer CJ, Price AL, Lettre G, Loos RJF, Weedon MN, Ingelsson E, O’Connell JR, Abecasis GR, Chasman DI, Goddard ME, Visscher PM, Hirschhorn JN, Frayling TM. 2014. Defining the role of common variation in the genomic and biological architecture of adult human height. Nat Genet 46:1173–1186. doi: 10.1038/ng.3097

Wray NR, Wijmenga C, Sullivan PF, Yang J, Visscher PM. 2018. Common Disease Is More Complex Than Implied by the Core Gene Omnigenic Model. Cell. doi: 10.1016/j.cell.2018.05.051

Wray NR, Yang J, Hayes BJ, Price AL, Goddard ME, Visscher PM. 2013. Pitfalls of predicting complex traits from SNPs. Nat Rev Genet 14:507–515. doi: 10.1038/nrg3457

Yang J, Benyamin B, McEvoy BP, Gordon S, Henders AK, Nyholt DR, Madden PA, Heath AC, Martin NG, Montgomery GW, Goddard ME, Visscher PM. 2010. Common SNPs explain a large proportion of the heritability for human height. Nat Genet 42:565–569. doi: 10.1038/ng.608

Yang J, Manolio TA, Pasquale LR, Boerwinkle E, Caporaso N, Cunningham JM, De Andrade M, Feenstra B, Feingold E, Hayes MG, Hill WG, Landi MT, Alonso A, Lettre G, Lin P, Ling H, Lowe W, Mathias RA, Melbye M, Pugh E, Cornelis MC, Weir BS, Goddard ME, Visscher PM. 2011. Genome partitioning of genetic variation for complex traits using common SNPs. Nat Genet 43:519–525. doi: 10.1038/ng.823

Zeh JA, Bonilla MM, Adrian AJ, Mesfin S, Zeh DW. 2012. From father to son: Transgenerational effect of tetracycline on sperm viability. Sci Rep 2:375. doi: 10.1038/srep00375

Zhang W, Reeves GR, Tautz D. 2020. Identification of a genetic network for an ecologically relevant behavioural phenotype in Drosophila melanogaster. Mol Ecol 29:502–518. doi: 10.1111/mec.15341

Zhou X, Stephens M. 2012. Genome-wide efficient mixed-model analysis for association studies. Nat Genet 44:821–824. doi: 10.1038/ng.2310

